# Experience-dependent modulation of the visual evoked potential: testing effect sizes, retention over time, and associations with age in 415 healthy individuals

**DOI:** 10.1101/2020.01.27.916692

**Authors:** Mathias Valstad, Torgeir Moberget, Daniël Roelfs, Nora B. Slapø, Clara M.F. Timpe, Dani Beck, Geneviève Richard, Linn Sofie Sæther, Beathe Haatveit, Knut Andre Skaug, Jan Egil Nordvik, Christoffer Hatlestad-Hall, Gaute T. Einevoll, Tuomo Mäki-Marttunen, Lars T. Westlye, Erik G. Jönsson, Ole A. Andreassen, Torbjørn Elvsåshagen

## Abstract

Experience-dependent modulation of the visual evoked potential (VEP) is a promising proxy measure of synaptic plasticity in the cerebral cortex. However, existing studies are limited by small to moderate sample sizes as well as by considerable variability in how VEP modulation is quantified. In the present study, we used a large sample (n = 415) of healthy volunteers to compare different quantifications of VEP modulation with regards to effect sizes and retention of the modulation effect over time. We observed significant modulation for VEP components C1 (Cohen’s *d* = 0.53), P1 (*d* = 0.66), N1 (*d* = −0.27), N1b (*d* = −0.66), but not P2 (p = 0.1), and in one time-frequency cluster (~30 Hz and ~70 ms post-stimulus; *d* = −0.48), 2-4 minutes after 2 Hz prolonged visual stimulation. For components N1 (*d* = −0.21) and N1b (*d* = −0.38), as well for the time-frequency cluster (*d* = −0.33), this effect was retained after 54-56 minutes. Moderate to high correlations (*ρ* = [0.39, 0.69]) between modulation at different postintervention blocks revealed a relatively high temporal stability in the modulation effect for each VEP component. However, different VEP components also showed markedly different temporal retention patterns. Finally, P1 modulation correlated positively with age (t = 5.26), and was larger for female participants (t = 3.91), with no effects of either age or sex on N1 and N1b potentiation. These results provide strong support for VEP modulation, and especially N1b modulation, as a robust measure of synaptic plasticity, but underscore the need to differentiate between components, and to control for demographic confounders.

## Introduction

Due to the essential role of synaptic plasticity in learning and memory (Takeuchi, Duszkiewicz, & Morris, 2013), as well as its likely role in the etiology of a range of psychiatric disorders (Consortium, 2014; Stephan, Baldeweg, & Friston, 2006), several non-invasive methodologies for studying long term potentiation (LTP)-like synaptic plasticity in humans have been developed. Among these approaches, the application of high frequency or prolonged visual stimulation to manipulate visual evoked potentials (VEPs) measured using electroencephalography (EEG) has proven especially promising (Cooke & Bear, 2012). Supporting the utility of this experimental paradigm in clinical research, modulation of VEP components after high frequency or prolonged visual stimulation appears to be altered in mood (Elvsåshagen et al., 2012; Normann, Schmitz, Fürmaier, Döing, & Bach, 2007) and psychotic illnesses (Çavuş et al., 2012). However, the specific VEP components exhibiting robust modulation effects and differences between patients and controls, as well as the retention of modulation effects, have varied between studies, highlighting a need for further characterization of VEP modulation induced by prolonged visual stimulation in a large sample of healthy individuals.

In a standard VEP modulation paradigm, subjects are exposed first to reversing checkerboard or grating stimuli which elicit VEPs, then to a prolonged (e.g. Normann et al., 2007) or high-frequency version (e.g. Teyler et al., 2005) of the same stimulus, and lastly, after some delay, to the initial stimulation again, which now typically evokes a slightly modulated visual potential. Importantly, the mechanisms underlying such experience-dependent VEP modulation seem to share many characteristics with LTP, thus having earned the placeholder epithet *LTP-like plasticity*. In mice, both NMDAR antagonists like CPP, and AMPAR insertion-inhibitor GluR1-CT prevent experience-dependent VEP modulation from occurring (Frenkel et al., 2006). Also, electrical stimulation-induced LTP at thalamocortical synapses in the primary visual cortex (V1) enhances visual evoked potentials and inhibits further experience-dependent VEP modulation (Cooke & Bear, 2012). In humans, the spatial frequency- and orientation-specific receptive fields of V1 neurons have been exploited to demonstrate a specificity of experience-dependent VEP modulation that is consistent with the synaptic specificity characteristic of LTP (McNair et al., 2006; Ross et al., 2008).

Although all published studies have reported experience-dependent VEP modulation, the exact components modulated and the duration of modulation have varied between experiments (Table 1). In humans, the VEP is characterized by components separated in time, voltage polarity, and likely neural generators, with the largely negative C1 probably originating in the primary visual cortex (Di Russo, Martínez, Sereno, Pitzalis, & Hillyard, 2002) and occurring at ~50-90 ms post-stimulus, the positive P1 at ~80-120 ms and the negative N1 at ~130-200 ms, both probably originating in striate and extrastriate areas (Di Russo et al., 2002), and the positive and likely very complex P2 at ~200-300 ms post-stimulus. While some researchers (McNair et al., 2006; Ross et al., 2008; Teyler et al., 2005) demonstrated modulation of the relatively late-occurring N1b component exclusively, others have demonstrated an effect that is earlier and more widespread, with modulation of the P1 and N1 components (Elvsåshagen et al., 2012), and even of the C1 component (Çavuş et al., 2012; Normann et al., 2007). However, in the two studies demonstrating C1 modulation, opposite directions of effect were observed. The duration of VEP modulation has also varied between studies. Among the studies measuring VEP within the time range of classical LTP, that is, at least 30 minutes (Lisman, 2017) after prolonged or high frequency visual stimulation, one demonstrated retention of the modulation (Teyler et al., 2005), while another did not (Ross et al., 2008). Thus, it is also unclear to which extent early (< 30 minutes after high frequency or prolonged stimulation) and late (> 30 minutes after high frequency or prolonged stimulation) VEP modulation are associated, such that early VEP modulation could be taken as indicative of late. While some of the observed differences may be attributable to variations in experiment characteristics such as the specific visual stimulus used (grating or checkerboard), as well as the duration and frequency of stimulation, heterogeneity of results between studies that are similar in these respects seems to implicate error variance.

**Table 1:** Overview of VEP modulation studies

| *Author, year<sup>a</sup>* | *N<sup>b</sup>* | *Intervention* | *Modulation<sup>c</sup>* |
| --- | --- | --- | --- |
| *Teyler et al., 2005* | 6 | 2 min 9 Hz checkerboard | N1b<sup>↓</sup> |
| *McNair et al., 2006* | 10 | 2 min 8.6 Hz grating | N1b<sup>↓</sup> |
| *Normann et al., 2007* | 32 | 10 min 2 Hz checkerboard | C1<sup>↑</sup>, P1<sup>↑</sup>, N1<sup>↓</sup> |
| *Ross et al., 2008* | 18 | 2 min 8.6 Hz grating | N1b<sup>↓</sup> |
| *Çavuş et al., 2012* | 41 | 2 min 8.87 Hz checkerboard | C1<sup>↓</sup>, N1b<sup>↓</sup> |
| *Elvsåshagen et al., 2012* | 66 | 10 min 2 Hz checkerboard | P1<sup>↑</sup>, N1<sup>↓</sup> |
| *Forsyth et al., 2015* | 65 | 2 min 8.87 Hz checkerboard | C1<sup>↑</sup>, P2<sup>↑</sup> |
| *de Gobbi-Porto et al., 2015* | 17 | 2 min 9 Hz checkerboard | N1b<sup>↓</sup> |
| *Klöppel et al., 2015* | 37 | 10 min 2 Hz checkerboard | C1<sup>↑</sup>, P1<sup>↑</sup> |
| *Smallwood et al., 2015* | 21 | 2 min 8.6 Hz grating | N1b<sup>↓</sup> |
| *Forsyth et al., 2017* | 45 | 2 min 8.87 Hz checkerboard | C1<sup>↑</sup>, P2<sup>↑</sup> |
| *Jahsan et al., 2017* | 64 | 2 min 8.87 Hz checkerboard | N1b<sup>↓</sup>, P2<sup>↑</sup> |
| *Spriggs et al., 2017* | 49 | 2 min 8.6 Hz grating | N1b<sup>↓</sup>, P2a<sup>↑</sup> |
| *Spriggs et al., 2018* | 40 | 2 min 9 Hz grating | C1<sup>↓</sup>, N1<sup>↑</sup>, P2<sup>↑</sup> |
| *Sumner et al., 2018* | 20 | 2 min 9 Hz grating | P2<sup>↑</sup> |
| *Zak et al., 2018* | 58 | 10 min 2 Hz checkerboard | C1<sup>↑</sup>, P1<sup>↑</sup>, N1<sup>↓</sup> |
| *Abuleil et al., 2019* | 47 | 2 min 9 Hz checkerboard | P1<sup>↓</sup>, N1b<sup>↓</sup> |
| *Spriggs et al., 2019* | 28 | 2 min 8.6 Hz grating | N1b<sup>↓</sup> |
| *Wynn et al., 2019* | 65 | 4 min of 10 Hz grating, on/off 5s | P1<sup>↓</sup>, N1b<sup>↑</sup>, P2<sup>↑</sup> |

Table of studies using high frequency or prolonged visual stimulation to manipulate visual evoked potentials in humans.<sup>a</sup> Details in references. <sup>b</sup> Results for some participants may have been reported in more than one paper. <sup>c</sup> Due to differing methods of analysis between studies, the exact nature of the modulated components can vary, and due to differences in statistical analysis between studies, the probability of actual modulation having been observed can also vary. Arrows denote direction of change pre-post intervention in the amplitude of a component (e.g. an upward arrow for a component that is negative at baseline means that the component became less negative or even positive after intervention).

Indeed, some of the studies at hand may have been underpowered with respect to differentiation between modulation of separate VEP components, and may not have controlled for adequate confounders. Potential confounders of the VEP modulation effect include the age and sex of participants. With age, there is a general decline in neural plasticity in animals (Burke & Barnes, 2006). Using the VEP modulation paradigm in humans, visual cortical plasticity has been demonstrated in older individuals in one sample (de Gobbi-Porto et al., 2015), but not in another (Spriggs, Cadwallader, Hamm, Tippett, & Kirk, 2017), and the relationship could be further elucidated with a continuous age distribution among participants. Further, sex differences in anatomical features such as cortical gyrification (Luders et al., 2004) might impact orientation of neural tissue, electrical conduction, and ultimately scalp EEG signals. Another factor that could impact observed VEP modulation is the level of attention afforded the visual stimulus, especially during high frequency or prolonged visual stimulation. Attention levels might be indexed by visual stimulation-driven steady state responses (Çavuş et al., 2012), or inversely by power in the alpha range (8-13 Hz) (Liu, Chiang, & Chu, 2013). The impact of such potential confounders should be further characterized to adequately evaluate effects of high frequency or prolonged visual stimulation in different populations.

There are multiple ways of quantifying VEP modulation, potentially leading to questionable analytic flexibility if the outcome is not defined a priori. For instance, while some researchers have focused on the N1b component of the VEP, which is typically operationalized as mean amplitude between the first negative and halfway to the first positive peak after P1 (e.g. McNair et al., 2006; Spriggs et al., 2017), others have focused on the N1 component, operationalized as the amplitude of the first negative peak after P1 (Elvsåshagen et al., 2012). Quantifications of VEP modulation to consider include changes from baseline to postintervention amplitudes in the C1, P1, N1, N1b, and P2 components, as well as in the peak to peak difference P1-N1. Furthermore, as the largest effects are not necessarily phase-locked, time-frequency analyses of the post-stimulus EEG should be employed to complement time-domain analyses. Since these components have not been directly compared in a large sample of healthy individuals, it is currently unknown which of the many potential indices of LTP-like synaptic plasticity is most sensitive and robust. Typical sample sizes within the field might make some studies vulnerable to winner’s curse and random effects (Ioannidis, 2008). Here, we conducted the largest study of VEP modulation to date in 415 healthy volunteers and directly compared several quantifications of VEP modulation, enabling us to obtain realistic effect sizes and to determine which quantifications are best suited for indexing LTP-like synaptic plasticity in humans.

The present study had three main aims: first, to determine which EEG measures exhibit robust modulation following prolonged visual stimulation; second, to assess the retention of such VEP modulation effects over intervals reaching the time range of LTP, and the correlations between the magnitude of early and late VEP modulation; and third, to examine the extent to which age, sex, and markers of attention might influence VEP modulation.

## Methods

### Participants

415 participants (age range: 18-88, 59% female) were recruited to this study from Statistics Norway and announcements in national news outlets, and included after screening for self-reported neurological or psychiatric disease. All participants had normal or corrected-to-normal vision. The experiment was approved by the Regional Ethical Committee of South-Eastern Norway, and all participants provided written informed consent.

### Experimental procedures

The VEP modulation paradigm was adopted from Normann et al. (2007). Over a period of 67 minutes, 11 VEP blocks, i.e., 2 baseline blocks, 1 intervention block of prolonged visual stimulation, and 8 postintervention blocks, were presented on a 24 inch 144Hz AOC LCD screen with 1 ms grey-to-grey response time (Fig. 1). All blocks, including the intervention block, consisted of a reversing checkerboard pattern with a spatial frequency of 1 cycle/degree over a ~28° visual angle. The reversal frequency was fixed at 2 reversals per second for the intervention block, whereas the baseline and postintervention blocks had jittered stimulus onset asynchronies of 500-1500 ms (mean = 1000 ms). All baseline and postintervention blocks lasted ~40 seconds (i.e., 40 checkerboard reversals), while the stimulation block lasted 10 minutes (i.e., 1200 reversals). Postintervention blocks were presented at 2 min, 3 min 40 s, 6 min 20 s, 8 min, ~30 min, ~32 min, ~54 min, and ~56 min after the intervention block. Through all blocks, the participants fixed their gaze on a red dot in the centre of the screen, and pressed a key on a gaming controller when its color changed from red to green. Between the seventh and eight, and between the ninth and tenth blocks, participants underwent mismatch negativity (Näätänen, Gaillard, & Mäntysalo, 1978) and prepulse inhibition (Graham & Murray, 1977) tasks, respectively.

**Figure 1.**
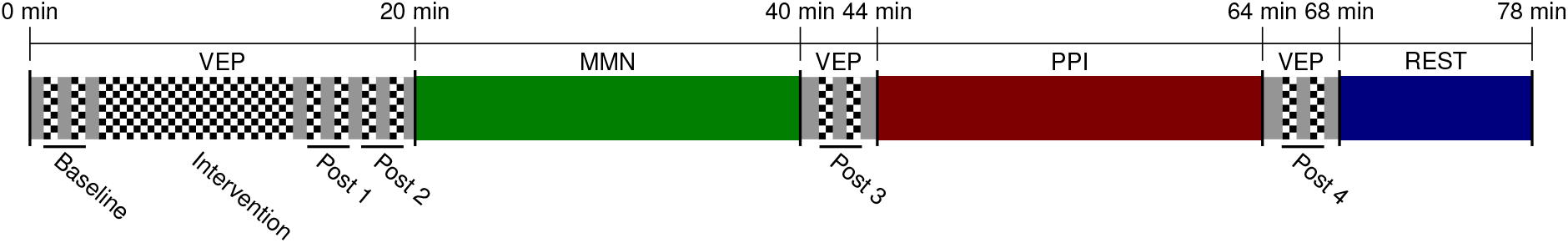
Experimental timeline. **VEP**: visual evoked potential paradigm, **MMN**: mismatch negativity paradigm, **PPI**: prepulse inhibition paradigm, **REST**: resting state EEG.

### Data acquisition

EEG recordings were acquired using a 64 channel BioSemi ActiveTwo amplifier, with Ag-AgCl sintered electrodes distributed across the scalp according to the international 10-20 system. External electrodes were placed at the outer canthi of both eyes (LO1, LO2), and below and above the left eye (IO1, SO1) in order to acquire horizontal and vertical electro-oculograms for eye movement and eye blink correction. Potentials at electrode sites were measured with respect to a common mode sense, with a driven right leg electrode minimizing common mode voltages, and sampled at 2048 Hz.

### Signal processing

Signal processing was performed using MATLAB and the EEGLAB toolbox for MATLAB (Delorme & Makeig, 2004), while statistical analysis was performed using R version 3.6.0 (R Core Team, 2019). Offline, files were downsampled to 512 Hz. Noisy channels were identified with PrepPipeline algorithms (Bigdely-Shamlo, Mullen, Kothe, Su, & Robbins, 2015) using default criteria, and removed. Remaining channels were first referenced to their common average voltage, before interpolation of removed channels from surrounding channel potentials, and finally all channels were rereferenced to the new common average after interpolation of bad channels. Data destined for time domain analysis were band-pass filtered between 0.1 and 40 Hz, while data for spectral analysis were high-pass filtered at 0.1 Hz. A fixed 20 ms delay in the visual presentation relative to the event markers was detected using a BioSemi PIN diode placed in front of the screen while running the paradigm, and event markers were adjusted offline to account for this. Next, epochs were extracted at 200 ms pre- to 500 ms post-stimulus. Muscle, eye blink and eye movement artifactual components were removed with SASICA defaults (Chaumon, Bishop, & Busch, 2015) after subjecting the epoched data to independent component analyses with the SOBI algorithm (Belouchrani, Abed-Meraim, Cardoso, & Moulines, 1993). Finally, epochs with amplitude diversions exceeding 100 μV were removed, and all channels were referenced to the AFz electrode.

### Data analysis

Three different modes of EEG analysis were pursued: time domain analyses at group and individual levels, frequency domain analyses at the individual level, and time-frequency analyses at group and individual levels. Since the baseline consisted of two VEP blocks, postintervention blocks were also collapsed into series of two blocks for equal comparison, resulting in one baseline assessment and four postintervention assessments.

For time domain analysis, C1 was defined as minimum amplitude between 50-100 ms post-stimulus, P1 as maximum amplitude between 80-140 ms, N1 as the amplitude of the first negative peak after P1, N1b as mean amplitude between the first negative and halfway to the first positive peak after P1 (effectively 150-190 ms post-stimulus), and P2 as mean amplitude in the 50 ms after and including the first positive peak after P1 (effectively 228-278 ms post-stimulus), reflecting increased latency variabilities with later components. C1 identification was quality controlled by visual inspection, and analyses were run with and without corrected data. In addition, we performed a completely data-driven, exploratory analysis, where voltages at each post-stimulus time point were calculated and assessed for postintervention changes. All channels were subjected to group-level time domain analysis, and the channel with highest amplitudes and most pronounced VEP modulation (i.e., Oz) was selected for all later analyses (Fig. 3).

For frequency domain analyses, entire continuous stretches of intervention block EEG were subjected to a Fast Fourier Transform (Cohen, 2014) before extraction of mean power within the alpha (8-13 Hz) and narrow steady state bands centered on the 2 Hz visual stimulation frequency (1.8-2.2 Hz).

For time-frequency analyses, high-pass filtered epochs from all participants were convolved with 5-cycle complex Morlet wavelets (Cohen, 2014) at each integer frequency between 10 and 120 Hz. To calculate induced power in addition to total power, each participant’s ERP at each assessment was also convolved with the same complex Morlet wavelets, and the resulting inner products were subtracted from the inner products at corresponding times and frequencies for each epoch. For both total and induced spectra, median amplitudes of inner products at each time point and each frequency were computed across assessments, and were decibel converted with a baseline between 150 and 100 ms pre-stimulus. The resulting pixels (representing a specific time-frequency combination) were then permuted across baseline and the first postintervention assessment in 2000 simulations, generating a null distribution for each pixel. The decibel values for each time point and frequency were then permuted again, simulation pixels were thresholded at p < 0.05 compared to the pixel null distributions, and the size of the largest resulting cluster was stored to generate a null distribution of cluster sizes. Finally, the actual decibel values were thresholded at p < 0.0005, and resulting clusters larger than 0.9995 of the null clusters were selected for further analysis. At the individual participant level, average power within the resulting clusters was then extracted at all assessments.

### Outcomes

Primary outcomes were i) modulation of components C1, P1, N1, N1b, and P2, as well as in the P1-N1 composite, between baseline and each postintervention assessment, ii) modulation of within time-frequency clusters total power between baseline and each postintervention assessment, and linear models for the effects of induced and evoked power for such differences, iii) correlations between baseline to postintervention amplitude changes for all components at all postintervention assessments, and iv) effects of age, gender, and steady-state and alpha band powers during prolonged visual stimulation on the subsequent modulation of components C1, P1, N1, N1b, and P2.

Raw values are reported along with standard errors, calculated as standard deviations over the square root of the sample size. Baseline to postintervention changes (i.e., modulation effects) are expressed as Cohen’s *d_z_* (henceforth denoted *d*), calculated as difference means over difference standard deviations (Cohen, 1988), and as response rates (rr), defined as the proportion of participants exhibiting the expected direction of baseline to postintervention changes. Correlations are expressed as Spearman’s *ρ*. P-values were calculated based on 20000 permutations between baseline and postintervention assessments, and are reported in their raw form. Alpha levels were adjusted to control for multiple comparisons according to the effective number of independent comparisons, derived using eigenvalues of the correlation matrix of the entire continuous data set (Li & Ji, 2005), yielding an experiment-wide significance threshold at 0.0009. Regression models were fitted using the general linear model, while controlling for baseline amplitudes, model fit is indexed using Nagelkerke R^2^, and effect is expressed with *t*-values.

## Results

The checkerboard reversal stimulation evoked the expected C1, P1, N1, and P2 components of the VEP (Fig. 2; see Table 2 for latencies and amplitudes). Initial group level analyses demonstrated that, across VEP components, the highest amplitudes and the largest modulation effects were exhibited at the occipital Oz electrode (Fig. 3A-B), which was accordingly selected for individual level analyses.

**Table 2:** VEP component amplitudes and latencies at baseline

| *Component* | *Latency (ms)* | *Amplitude (μV)* |
| --- | --- | --- |
| *C1* | 66.6±0.51 | -3.91±0.24 |
| *P1* | 99.0±0.41 | 8.42±0.30 |
| *N1* | 140.3±0.81 | -5.92±0.24 |
| *N1b* | NA | -1.65±0.20 |
| *P2* | NA | 1.41±0.17 |

Table of VEP component amplitudes and latencies at baseline, measured at the occiput (Oz) with anterior reference (AFz). **NA**: not applicable.

**Figure 2.**
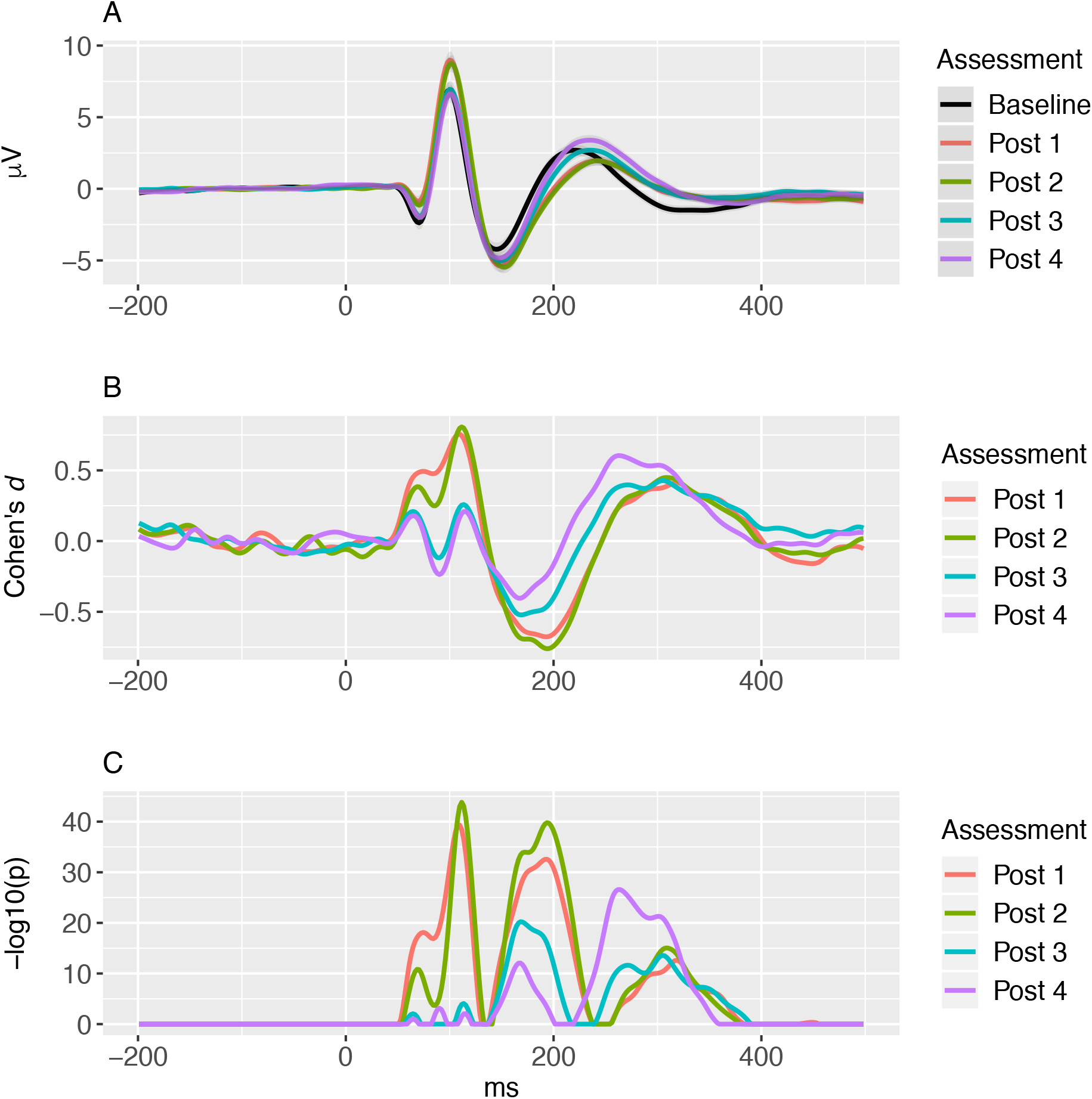
**A.** Grand average visual evoked potentials measured at the occiput (Oz) with anterior reference (AFz) at baseline, post 1 (2-4 min after prolonged visual stimulation), post 2 (6-8 min), post 3 (30-32 min), and post 4 (54-56 min). **B.** Cohen’s *d* from baseline VEP and the postintervention assessments. **C.** P-values for difference between baseline VEP the postintervention assessments, Bonferroni corrected and log transformed for visualization purposes.

**Figure 3.**
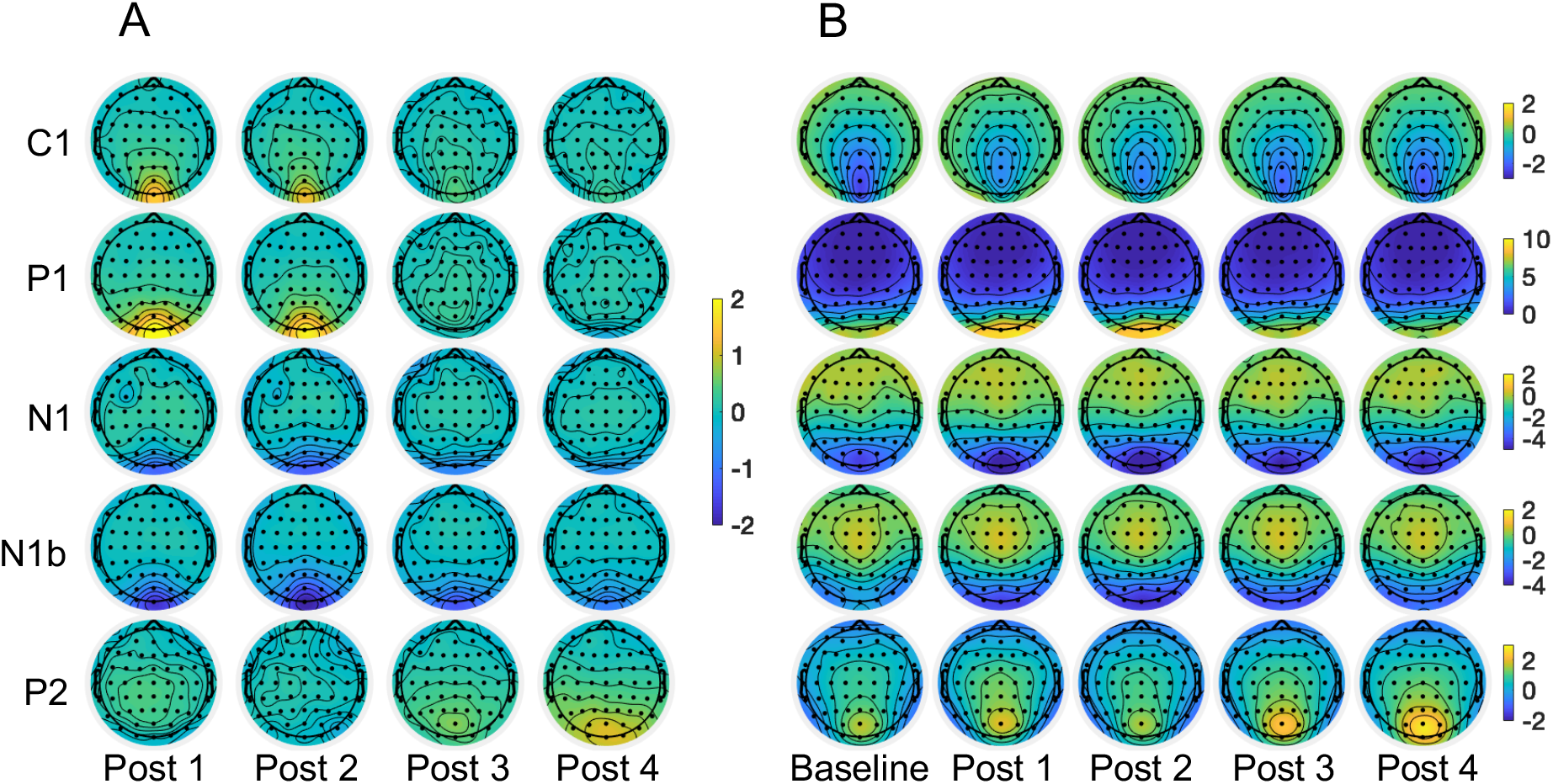
**A.** Scalp topographical distribution of C1, P1, N1, N1b, and P2 unscaled amplitude differences (in μV) from baseline to postintervention assessments 1 (2-4 min after prolonged visual stimulation), 2 (6-8 min), 3 (30-32 min), and 4 (54-56 min). **B.** Scalp topographical distribution of C1, P1, N1, N1b, and P2 amplitudes at baseline and each of the postintervention assessments 1-4.

When testing for modulation effects across all timepoints of the VEP at the first postintervention assessment after prolonged visual stimulation, significant changes at latencies of 54.7-128.9 ms, 138.7-234.4 ms, and 257.8-375.0 ms were observed (Fig. 2). Correspondingly, experience-dependent VEP modulation was apparent as amplitude changes from baseline to the first postintervention assessment for both the C1 (*d* = 0.53, rr = 0.70), P1 (*d* = 0.66, rr = 0.76), N1 (*d* = −0.27, rr = 0.62), N1b (*d* = −0.66, rr = 0.77), but not P2 (p = 0.1, rr = 0.53) components, with highly similar effects for both the C1 (*d* = 0.43, rr = 0.67), P1 (*d* = 0.55, rr = 0.72), N1 (*d* = −0.26, rr = 0.61), N1b (*d* = −0.71, rr = 0.77) and the P2 (p = 0.1, rr = 0.54) components at the immediately following second postintervention assessment. Some, but not all, changes after prolonged visual stimulation were retained at the third and fourth postintervention assessments. The C1 component retained modulation at the third (*d* = 0.20, rr = 0.58), and tendentially at the fourth (*d* = 0.16, p = 0.001, rr = 0.56) postintervention assessment. The P1 component did not retain modulation at the third (p = 0.38, rr = 0.54), nor at the fourth (p = 0.22, rr = 0.48) postintervention assessment. The N1 component retained modulation at the third (*d* = −0.17, rr = 0.60), and fourth (*d* = −0.21, rr = 0.66) postintervention assessment. The N1b component retained modulation at both the third (*d* = −0.53, rr = 0.75), and the fourth (*d* = −0.38, rr = 0.68) postintervention assessment. Finally, the P2 component gained modulation at the third (*d* = 0.30, rr = 0.65) and the fourth (*d* = 0.54, rr = 0.75) postintervention assessment (Table 3, Fig. 4).

**Table 3:** VEP component modulation after prolonged visual stimulation

|  |  | *C1* | *P1* | *N1* | *N1b* | *P2* |
| --- | --- | --- | --- | --- | --- | --- |
| *Post 1* | *d* | 0.53 | 0.66 | -0.27 | -0.66 | 0.08 |
| (2-4 | *rr* | 0.70 | 0.76 | 0.62 | 0.77 | 0.53 |
| *min*) | *p* | $<5*10^{-5}$ | $<5*10^{-5}$ | $<5*10^{-5}$ | $<5*10^{-5}$ | 0.10 |
| | $-\log(p)$ | 23.1 | 33.7 | 7.1 | 33.7 | 1 |
| *Post 2* | *d* | 0.44 | 0.55 | -0.26 | -0.71 | 0.08 |
| (6-8 | *rr* | 0.67 | 0.72 | 0.61 | 0.77 | 0.54 |
| *min*) | *p* | $<5*10^{-5}$ | $<5*10^{-5}$ | $<5*10^{-5}$ | $<5*10^{-5}$ | 0.10 |
| | $-\log(p)$ | 16.9 | 24.6 | 6.8 | 37.8 | 1 |
| *Post 3* | *d* | 0.20 | 0.04 | -0.17 | -0.53 | 0.30 |
| (30-32 | *rr* | 0.58 | 0.54 | 0.60 | 0.75 | 0.65 |
| *min*) | *p* | $<5*10^{-5}$ | 0.38 | 0.0003 | $<5*10^{-5}$ | $<5*10^{-5}$ |
| | $-\log(p)$ | 4.16 | 0.4 | 3.2 | 23.0 | 8.9 |
| *Post 4* | *d* | 0.16 | -0.06 | -0.21 | -0.38 | 0.54 |
| (54-56 | *rr* | 0.56 | 0.48 | 0.66 | 0.68 | 0.75 |
| *min*) | *p* | 0.001 | 0.22 | $<5*10^{-5}$ | $<5*10^{-5}$ | $<5*10^{-5}$ |
| | $-\log(p)$ | 2.9 | 0.7 | 4.8 | 12.9 | 24.3 |

Table of VEP component modulation after prolonged visual stimulation. **d**: Cohen's *d*, **rr**: response rate, **p**: p-value after 20000 permutations, **$-\log(p)$** : negative decimal logarithm of t-test p-value (for illustration, not all modulations are normally distributed).

**Figure 4.**
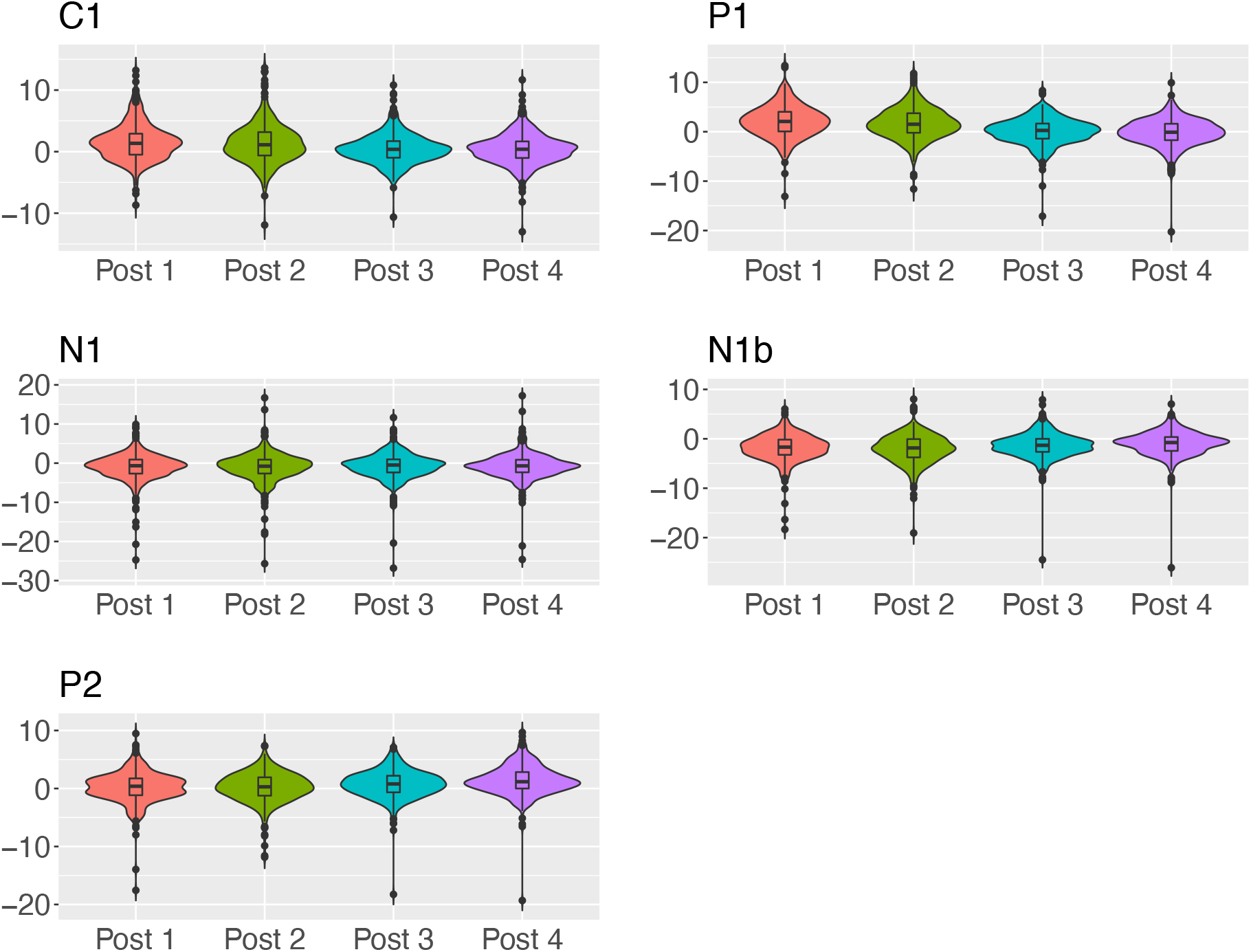
Distributions of amplitude differences (in μV) between baseline and postintervention assessments post 1 (2-4 min after prolonged visual stimulation), post 2 (6-8 min), post 3 (30-32 min), and post 4 (54-56 min), for VEP components C1, P1, N1, N1b, and P2 (n = 415).

The P1-N1 composite exhibited significant modulation at the first (*d* = 0.70, rr = 0.80), second (*d* = 0.60, rr = 0.78), and third (*d* = 0.19, rr = 0.61), but not the last (*d* = 0.14, rr = 0.60) postintervention assessment.

There were also differences between component amplitudes within assessments (Fig. 5), with significant changes from the first to the second baseline block for components P1 (*d* = 0.21), N1 (*d* = −0.39), N1b (*d* = −0.28), and P2 (*d* = 0.17), from the first to the second postintervention block for components C1 (*d* = 0.18), N1 (*d* = 0.24), and from the seventh to the eighth postintervention block for components C1 (*d* = 0.24), P1 (*d* = 0.31), N1 (*d* = −0.19), and N1b (*d* = −0.35). These effects were weaker than effects of the prolonged visual stimulation for components C1 (p = 1.2 × 10^−9^), P1 (p = 4.3 × 10^−14^), N1 (p = 2.4 × 10^−4^), and N1b (p= 1.1 × 10^−15^), but not P2 (p = 0.66).

**Figure 5.**
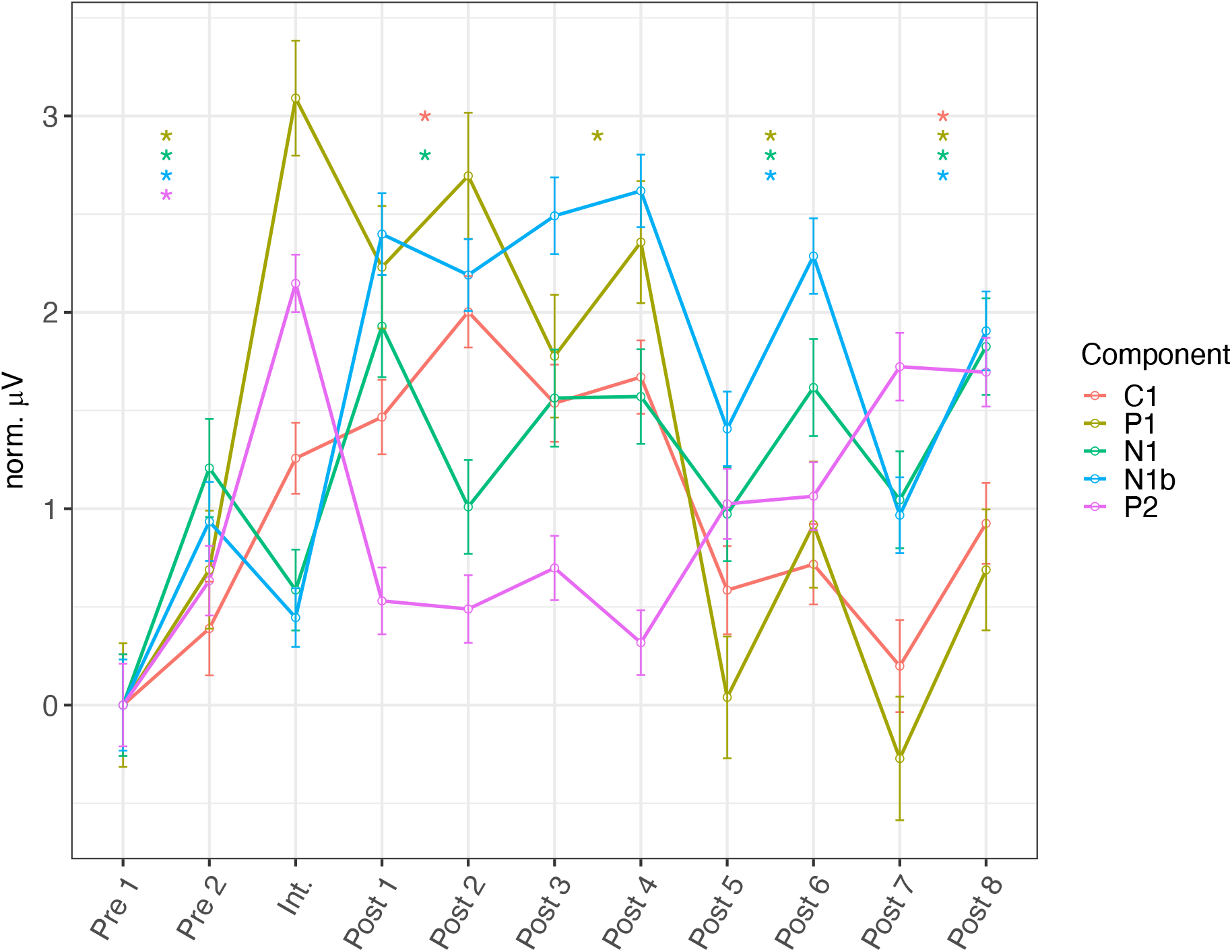
Component amplitudes at separate checkerboard stimulation blocks, normalized to the first block, with error bars showing standard error of measurement. Asterisks denote significant (p < 0.0009) amplitude change within assessments (i.e. from pre 1 to pre 2, from post 1 to post 2, from post 3 to post 4, from post 5 to post 6, and from post 7 to post 8). **Int.**: Intervention block.

The time-frequency analysis exploring the main effect of prolonged visual stimulation yielded five significant clusters (Fig. 6). Results from analyses across assessments using individual participants’ values averaged within clusters are presented in Table 4. Notably, these revealed that only the first cluster exhibited modulation at all postintervention assessments, including the first (*d* = −0.48, rr = 0.65), second (*d* = −0.60, r = 0.72), third (*d* = −0.44, rr = 0.66), and fourth (*d* = −0.33, rr = 0.65). This cluster was centered around ~30 Hz and ~70 ms post-stimulus, and the power reduction after prolonged visual stimulation was well modeled (R^2^ = 0.31) by power changes in a corresponding induced cluster (t = 7.22, p = 2.7 × 10^−12^), C1 modulation (t = −6.57, p = 1.7 × 10^−10^), and P1 modulation (t = 6.43, p = 3.8 × 10^−10^).

**Table 4:** Cluster power modulation after prolonged visual stimulation

|  |  | *A1* | *A2* | *A3* | *A4* | *A5* | *I1* |
| --- | --- | --- | --- | --- | --- | --- | --- |
| *Post 1* | *d* | -0.48 | 0.19 | -0.19 | 0.17 | -0.16 | -0.34 |
| (2-4 | *rr* | 0.65 | 0.56 | 0.58 | 0.59 | 0.54 | 0.62 |
| *min*) | *p* | $<5*10^{-5}$ | 0.0001 | 0.0001 | 0.0004 | 0.001 | $<5*10^{-5}$ |
| | $-\log(p^*)$ | 19.9 | 4.1 | 4.2 | 3.4 | 3.0 | 11.0 |
| *Post 2* | *d* | -0.60 | 0.35 | -0.09 | 0.06 | -0.16 | -0.41 |
| (6-8 | *rr* | 0.72 | 0.62 | 0.53 | 0.53 | 0.56 | 0.65 |
| *min*) | *p* | $<5*10^{-5}$ | $<5*10^{-5}$ | 0.06 | 0.23 | 0.0001 | $<5*10^{-5}$ |
| | $-\log(p^*)$ | 28.5 | 11.1 | 1.2 | 0.64 | 2.9 | 15.2 |
| *Post 3* | *d* | -0.44 | 0.09 | -0.20 | -0.36 | -0.12 | -0.30 |
| (30-32 | *rr* | 0.66 | 0.53 | 0.55 | 0.65 | 0.55 | 0.61 |
| *min*) | *p* | $<5*10^{-5}$ | 0.056 | $5*10^{-5}$ | $<5*10^{-5}$ | 0.015 | $<5*10^{-5}$ |
| | $-\log(p^*)$ | 17.2 | 1.3 | 4.1 | 11.7 | 1.8 | 8.7 |
| *Post 4* | *d* | -0.33 | 0.15 | -0.18 | -0.28 | -0.17 | -0.18 |
| (54-56 | *rr* | 0.65 | 0.57 | 0.59 | 0.62 | 0.55 | 0.57 |
| *min*) | *p* | $<5*10^{-5}$ | 0.003 | 0.0001 | $<5*10^{-5}$ | 0.001 | 0.0004 |
| | $-\log(p^*)$ | 10.4 | 2.5 | 3.7 | 7.8 | 3.1 | 3.4 |

Table of cluster power modulations after prolonged visual stimulation. **d**: Cohen's *d*, **rr**: response rate, **p**: p-value after 20000 permutations, **$-\log(p)$** : negative decimal logarithm of t-test p-value (for illustration, not all potentiations are normally distributed), **A1-5**: Cluster absolute power, **I1**: Induced power in the first cluster.

**Figure 6.**
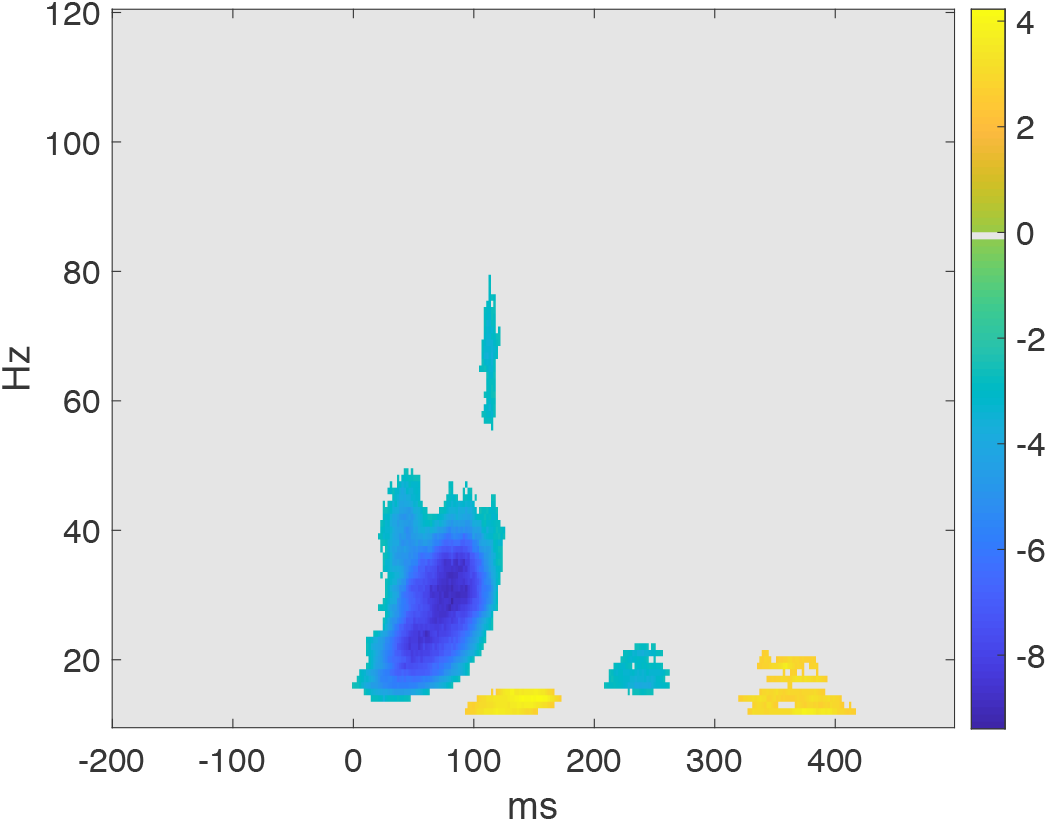
Changes in total power in frequencies 10-120, before to after prolonged visual stimulation, given as t-scores for each pixel, within significant clusters.

Correlations across assessments for baseline to postintervention modulation effects were moderate, ranging from Spearman’s *ρ* = [0.47, 0.69] for C1, *ρ* = [0.39, 0.67] for P1, *ρ* = [0.42, 0.62] for N1, *ρ* = [0.44, 0.66] for N1b, and *ρ* = [0.47, 0.60] for the P2 component (Fig. 7). All correlations above and including r = 0.17 remained significant after multiple comparison correction.

**Figure 7.**
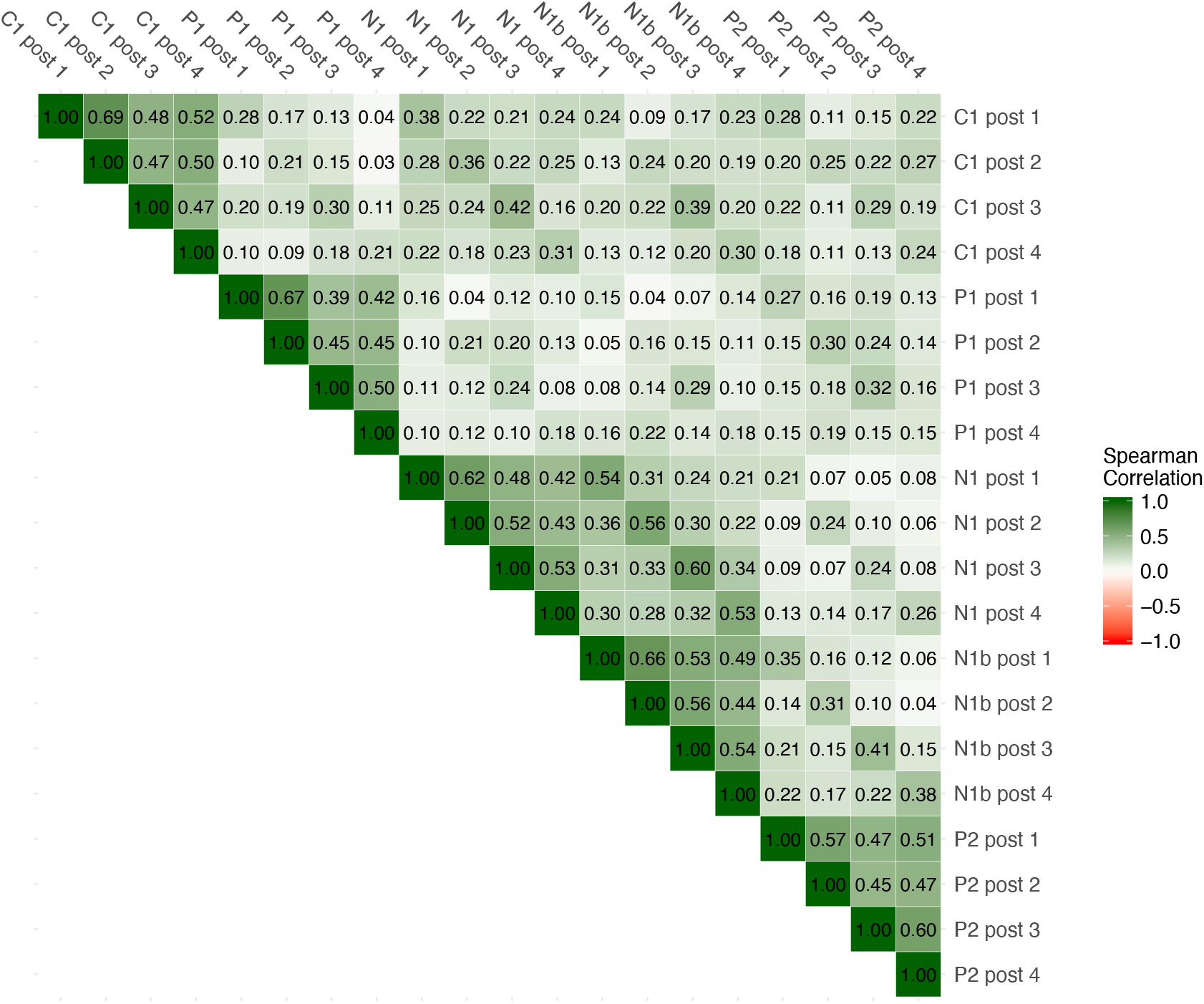
Spearman’s *ρ* correlations between modulations of VEP components C1, P1, N1, N1b, and P2 at postintervention assessments 1-4.

The regression model for P1 modulation (R^2^ = 0.15), revealed effects of age (t = 5.26, p = 1.6 × 10^−7^) and sex (t = 3.91, p = 9.7 × 10^−5^), with greater modulation for older participants and female participants, respectively. The regression model for P2 modulation (R^2^ = 0.09) also showed an increased difference from baseline to postintervention blocks for female participants (t = 5.08, p = 4.3 × 10^−7^). The regression model for C1 modulation (R^2^ = 0.11) revealed an interaction effect of age and time (t = 4.35, p = 1.5 × 10^−5^), indicating that while the postintervention modulation for younger participants vane throughout the experiment, this is less the case for older participants. The regression model for the major time-frequency component (R^2^ = 0.03) revealed an effect of age (t = −4.56, p = 5.7 × 10^−6^). Regression models for N1 (R^2^ = 0.04) and N1b (R^2^ = 0.07) modulation did not provide evidence for effects of age, sex, intervention block alpha power, or intervention steady state power. Finally, for the attentional task, we only obtained hit rate data for 45.8% of participants, due to error in the gaming controller. Thus, we performed a set of control analyses to ensure that the participants for which attentional data was not obtained did not differ from the participants for which attentional data was obtained. These showed that there was no difference between these groups in P1, N1, N1b, or P2 modulation, but only a nominal difference in C1 modulation (p = 0.04), and that clear VEPs were evoked for 96% of participants for which attentional data was not obtained. Among participants for which attentional data was obtained, the mean hit rate was 98.4%. Together, these results indicate overall satisfying levels of attention.

## Discussion

The current study yielded four main findings. First, we demonstrate robust experience-dependent modulation of the visual evoked potential in a large sample of healthy volunteers (n = 415). Second, the retention of this modulation effect over time varied across VEP components, strongly suggesting that VEP modulation is not a unitary phenomenon and likely involves several different plasticity mechanisms. Third, age and sex emerged as significantly associated with some, but not all, quantifications of VEP modulation, while electrophysiological indices of attention appeared unrelated to the degree of modulation. Finally, we identify the N1b component as the most sensitive quantification of both early (2-4 min post-intervention) and late (54-56 min post-intervention) VEP modulation.

### Experience-dependent modulation of visual evoked potentials

At the first and second postintervention assessments, respectively 2 and 6 minutes after prolonged visual stimulation, moderate to strong modulation was observed in VEP components C1, P1, N1, and N1b, as well as in the composite P1-N1. Such experience-dependent modulations have previously been shown to share many characteristics with LTP, such as NMDAR-dependence (Frenkel et al., 2006), post-synaptic AMPAR insertion dependence (Frenkel et al., 2006), and stimulus specificity (McNair et al., 2006; Ross et al., 2008), and have therefore been regarded as indices of LTP-like cortical synaptic plasticity. We have shown that the quantifications of VEP modulation that have previously been described in the literature – modulations of the C1, P1, N1, and N1b components – coincide with the latencies at which the post-stimulus VEP exhibited modulation after prolonged visual stimulation in the present study.

Time-frequency analyses also revealed differences in total power at several latencies and frequencies, of which only one cluster (~70 ms and ~30 Hz) exhibited effects of prolonged visual stimulation that were comparable to effects seen on time domain VEP components. Since these time-frequency modulations were independent of time domain VEP modulations at comparable latencies, they might reflect neural dynamics to which time domain VEP modulations are not sensitive.

### Experience-dependent VEP modulation: retention slopes and correlations

We observed differential response patterns between quantifications of VEP modulation, indicating differences in underlying mechanisms. Retention at the third and fourth postintervention assessments, i.e., ~30-32 and ~54-56 minutes after prolonged visual stimulation, was observed for components C1, N1, and N1b. The retention of C1, N1 and N1b modulation at 30 and 54 minutes postintervention is consistent with LTP-like synaptic processes as underlying mechanisms, since this duration goes beyond the usual decay of presynaptic short-term potentiation (Citri & Malenka, 2008; Regehr, 2012). Spearman correlations around 0.42-0.52 between C1, N1, and N1b modulations at 2 and 54-56 minutes postintervention suggest a connection between early and later modulation effects, which has been established for most forms of synaptic plasticity (Citri & Malenka, 2008), further corroborating the claim that C1, N1, and N1b modulations reflect LTP-like cortical plasticity.

With a sharp voltage increase in the intervention block and subsequent return to near baseline in the first two postintervention assessments, and renewed amplitude increases in the third and last postintervention assessments (Fig. 4; Fig. 5), the response pattern for the P2 component, similar to what has been observed previously (Forsyth, Bachman, Mathalon, Roach, & Asarnow, 2015; Forsyth et al., 2017), constitutes a clear exception, and appears inconsistent with NMDAR-dependent LTP, which exhibits a gradual decay (Citri & Malenka, 2008). Along the same lines, the P2 component appears to lack input specificity (Sumner et al., 2018). Thus, the effect of time on P2 amplitudes might seem to require some other mechanism than LTP-like synaptic plasticity. On the other hand, the retention slope of P1 is consistent with synaptic plasticity as underlying mechanism, although with a complete decay between 6 and 30 minutes after prolonged visual stimulation, P1 modulation might reflect some short-term plasticity such as post-tetanic potentiation (Citri & Malenka, 2008).

### Age and sex modulation of some, but not all, VEP components

Linear regression showed a positive main effect of age on P1 modulation, and a positive interaction effect between age and time after intervention for C1 modulation, but no effects of age on modulation of either the N1, N1b or the P2 components. These results are in line with a previous demonstration of robust VEP modulation among older individuals (de Gobbi-Porto et al., 2015), but seem to contrast with the lack of N1b modulation previously observed in older participants (Spriggs et al., 2017), and with the more general decline in neural plasticity associated with aging (Burke & Barnes, 2006). Further, regression models demonstrated larger P1 modulation, and larger increase in P2 amplitudes, among female participants, a result that – like the effects of age – was independent of baseline amplitudes. Together, these results underscore the need to differentiate between VEP components, and to control for demographic variables like age and sex, especially in case-control studies of VEP modulation.

Linear regression models for the effects of age, sex, intervention block alpha power and steady state power on the modulation of components C1, P1, N1, N1b, and P2 revealed no effects of attentional proxies on any of the quantifications of VEP modulation, suggesting that participants were sufficiently attentive to the prolonged visual stimulation for VEP modulation to occur. However, in a previous study of VEP modulation using 8.7 Hz visual stimulation (Çavuş et al., 2012), intervention block steady state power was associated with N1b modulation in healthy controls. Although neural entrainment to visual flickering can occur at frequencies between 1 and at least 50 Hz, the sensitivity at frequencies around the alpha band is higher than at 2 Hz (Herrmann, 2001), such that our 2 Hz prolonged visual stimulation may have been too slow for significant entrainment to occur.

### Robust and enduring modulation of component N1b

Our quantifications of VEP modulation seem to be relatively specific in that they exhibit distinct effects, retention slopes and associations with age and sex. Modulation of the N1b component after prolonged visual stimulation was overall the strongest effect. Effect size differences, relatively high correlations, and comparable associations with age and sex between components N1 and N1b suggest that N1b operationalizations might be preferable, at least under conditions similar to those present in this study. Although some observed effects of time might have been caused by other experimental characteristics than the prolonged visual stimulation, the N1b component has repeatedly been shown to increase in amplitude with high frequency visual stimulation, and not without (Teyler et al., 2005), and not with visual stimulation of a different orientation (Ross et al., 2008) or spatial frequency (McNair et al., 2006), supporting the notion that at least N1b modulation is due to the high frequency or prolonged visual stimulation.

### Possible influence of postintervention blocks on retention

In the present study we observed modulation of components P1, N1, N1b, and P2 even between blocks of short duration checkerboard stimulation. Thus, there is reason to question whether the retention, especially for components N1 and N1b which exhibit long duration modulation, could have been increased by the postintervention stimulus blocks. Postintervention blocks have been shown to decrease retention of N1b modulation (Teyler et al., 2005), but with frequency differences between intervention and postintervention blocks that were greater than in the present study, so some influence in favor of retention cannot be ruled out with the present data.

### Conclusion

The results of the current study show robust modulation after prolonged visual stimulation of VEP components C1, P1, N1, and N1b, as well as of ~30 Hz power at ~70 ms post-stimulus. Moreover, we observed differential retention slopes, effect sizes, and associations to age and sex for the modulation of VEP components, strongly suggesting that VEP modulation is not a unitary phenomenon. Taken together with results from a series of invasive studies in rodents, our current results support the use of prolonged visual stimulation induced VEP modulation, and especially N1b modulation, as a robust, non-invasive index of LTP-like cortical plasticity in humans.

## Acknowledgments

This study was funded by the Research Council of Norway, the South-Eastern Norway Regional Health Authority, Oslo University Hospital and a research grant from Mrs. Throne-Holst. The authors report no biomedical financial interests or potential conflicts of interest.

